# Forty high-intensity interval training sessions blunt exercise-induced changes in the nuclear protein content of PGC-1α and p53 in human skeletal muscle

**DOI:** 10.1101/580373

**Authors:** Cesare Granata, Rodrigo S.F. Oliveira, Jonathan P. Little, David J. Bishop

**Author notes:** **Corresponding author:** Cesare Granata, Department of Diabetes, Central Clinical School, Faculty of Medicine, Nursing and Health Sciences, Monash University, Alfred Centre, 99 Commercial Rd, Melbourne, 3004, VIC, Australia.

## Abstract

Exercise-induced increases in peroxisome proliferator-activated receptor γ coactivator-1α (PGC-1α) and p53 protein content in the nucleus mediate the initial phase of exercise-induced mitochondrial biogenesis. Here we investigated if exercise-induced increases in these and other markers of mitochondrial biogenesis were altered after 40 sessions of twice-daily high-volume high-intensity interval training (HVT) in human skeletal muscle. Vastus lateralis muscle biopsies were collected from 10 healthy recreationally active participants before, immediately post, and 3h after a session of HIIE performed at the same absolute exercise intensity before and after HVT (Pre-HVT and Post-HVT, respectively). The protein content of common markers of exercise-induced mitochondrial biogenesis were assessed in nuclear- and cytosolic-enriched fractions by immunoblotting; mRNA contents of key transcription factors and mitochondrial genes were assessed by qPCR. Despite exercise-induced increases in PGC-1α, p53, and plant homeodomain finger-containing protein 20 (PHF20) protein content, the phosphorylation of p53 and acetyl-CoA carboxylase (p-p53^Ser15^ and p-ACC^Ser79^, respectively), and PGC-1α mRNA Pre-HVT, no significant changes were observed Post-HVT. Forty sessions of twice-daily high-intensity interval training blunted all of the measured exercise-induced molecular events associated with mitochondrial biogenesis that were observed Pre-HVT. Future studies should determine if this loss relates to the decrease in relative exercise intensity, habituation to the same exercise stimulus, or a combination of both.

## Introduction

Mitochondria are responsible for the production of the majority of the energy required to sustain daily activities and are a key regulator of energy homeostasis (31). The importance of mitochondria is underlined by the links between a healthy mitochondrial pool and enhanced endurance performance (37), improved health (52), and a reduced risk of several lifestyle-related chronic diseases (6, 46). Exercise has long been known to induce mitochondrial biogenesis (34) - the making of new components of the mitochondrial reticulum (22). These adaptations to exercise training have been proposed to result from the cumulative effect of transient changes in nuclear protein content (73) and mRNA expression (55) induced by each exercise session.

Peroxisome proliferator-activated receptor γ coactivator-1α (PGC-1α) is a key regulator of exercise-induced mitochondrial biogenesis (74) (for an in-depth analysis of the effects of exercise on mitochondrial biogenesis mediated by PGC-1α [and p53] the reader is referred to some excellent reviews; (17, 21, 35, 64)). In both rat (73) and human (24, 33, 43, 44) skeletal muscle, it has been observed that there is a post-exercise increase of PGC-1α protein content in the nucleus, where PGC-1α performs its transcriptional activity (62). Changes in PGC-1α protein content (30), as well as the content of other proteins (e.g., p53 (36)), contribute to the exercise-induced upregulation of PGC-1α mRNA (58). Exercise-induced increases in the mRNA levels of PGC-1α and other genes (49, 55, 69), as well as the protein content of selected PGC-1α upstream regulators (48) and selected mitochondrial proteins and transcription factors (55, 69) measured in whole muscle lysates, have been shown to be reduced as a training intervention progresses. However, no study has investigated exercise-induced changes in the nuclear content of PGC-1α, or other important proteins modulating mitochondrial biogenesis, before and after a training intervention. Given that increased PGC-1α protein content in the nucleus represents an important process that contributes to the initial phase of exercise-induced mitochondrial biogenesis (73), it is important to better understand how the response of this transcriptional cofactor changes with training.

p53 is another important regulator of exercise-induced mitochondrial biogenesis in human skeletal muscle (64). Nuclear accumulation of p53 protein has been reported immediately (24), or 3 hours (70), after a single session of exercise. While the mechanisms underlying the nuclear accumulation of p53 are complex (47, 53), they have partly been attributed to phosphorylation of p53 at serine 15 (p-p53^Ser15^) (53) - a posttranslational modification that enhances p53 protein stability (68) and prevents its nuclear export and cytosolic degradation (32, 53). However, once again, these molecular events have only been investigated following a single exercise session, and it is not known if they are altered by training. Given that the majority of the p53 activity takes place in the nucleus (53), it is important to determine if the early events of the p53-mediated exercise-induced mitochondrial biogenesis are differentially regulated in this subcellular compartment as the training intervention progresses.

Therefore, the aim of our study was to investigate if a session of high-intensity interval exercise (HIIE), performed at the same absolute workload before and after a period of high-volume training (HVT; 40 sessions of high-intensity interval training [HIIT] performed twice-daily for 20 consecutive days), induces similar increases in the protein content of PGC-1α, p53, and p-p53^Ser15^ in the nucleus. Upstream signaling, as well as the mRNA content of several genes involved in exercise-induced mitochondrial biogenesis, were also investigated before and after HVT. The same absolute workload was chosen as this approach is often used in training studies (22), as well as in practice, where individuals regularly repeat the same exercise session. We hypothesized that 40 sessions of HIIT would result in significantly reduced exercise-induced increases in these events mediating exercise-induced mitochondrial biogenesis. Despite debate regarding how well exercise-induced molecular events can predict training-induced adaptations (22), findings from the present study will provide a better understanding of how molecular signals are altered when the same exercise stimulus is repeated. This will also improve our knowledge of the mechanisms underlying the common observation of smaller fitness gains as training progresses (42, 45), and may inform strategies to maintain the effectiveness of exercise to stimulate mitochondrial biogenesis.

## Materials and methods

### Participants

Ten healthy men (20 ± 2 y; 180 ± 12 cm; 80 ± 15 kg; 46.2 ± 7.6 mL · min^-1^ · kg^-1^), who were not regularly engaged in cycling-based sports, were moderately-trained (i.e., undertaking moderate, unstructured aerobic activity for less than 3 to 4 hours per week for at least 6 months prior to the study), and were non-smokers and free of medications, volunteered to participate in this study. Upon passing an initial medical screening participants were informed of the study requirements, risks, and benefits, before giving written informed consent. All experimental protocols and study procedures were approved by the Victoria University Human Research Ethics Committee and conformed to the standards set by the latest revision of the Declaration of Helsinki. All participants completed the study; however, due to the limited amount of muscle tissue harvested during the second biopsy trial, data from one participant were excluded (including physiological and performance data).

### Study design and testing

This research was part of a larger, previously-published study investigating the effect of different training volumes on mitochondrial adaptations (23). The experimental protocol specific to the portion of the study described in this manuscript consisted of three tests, each separated by 48 to 72 hours, repeated before and after the HVT: a 20-km cycling time trial (20k-TT), a graded exercise test (GXT) and a HIIE biopsy trial (Pre-HVT and Post-HVT). During the 20 days of HVT participants performed HIIT twice a day (Figure 1). Prior to beginning this phase of the larger study, participants were familiarized with the 20k-TT, the GXT and the HIIE, and completed the normal volume training (NVT) phase (12 HIIT sessions in 4 weeks; Figure 1). It has been reported that the transcriptional response to the first session of exercise (first bout effect (4)) can differ significantly from the response to subsequent exercise sessions (4, 50, 55, 72). Thus, the NVT phase served not only to habituate participants to the rigors of twice-daily HIIT during the HVT phase, but also to eliminate possible biases brought about by the “first-bout” effect (4). Finally, participants were required to refrain from vigorous exercise for the 72 h preceding each test, from alcohol for 24 h before testing, and from food and caffeine consumption for 3 h before each test. Although the lack of a “no exercise” control group could be considered a limitation of this study, it has previously been reported that there are no changes in upstream regulators of mitochondrial biogenesis in a “no exercise” control group (26).

**Figure 1.**
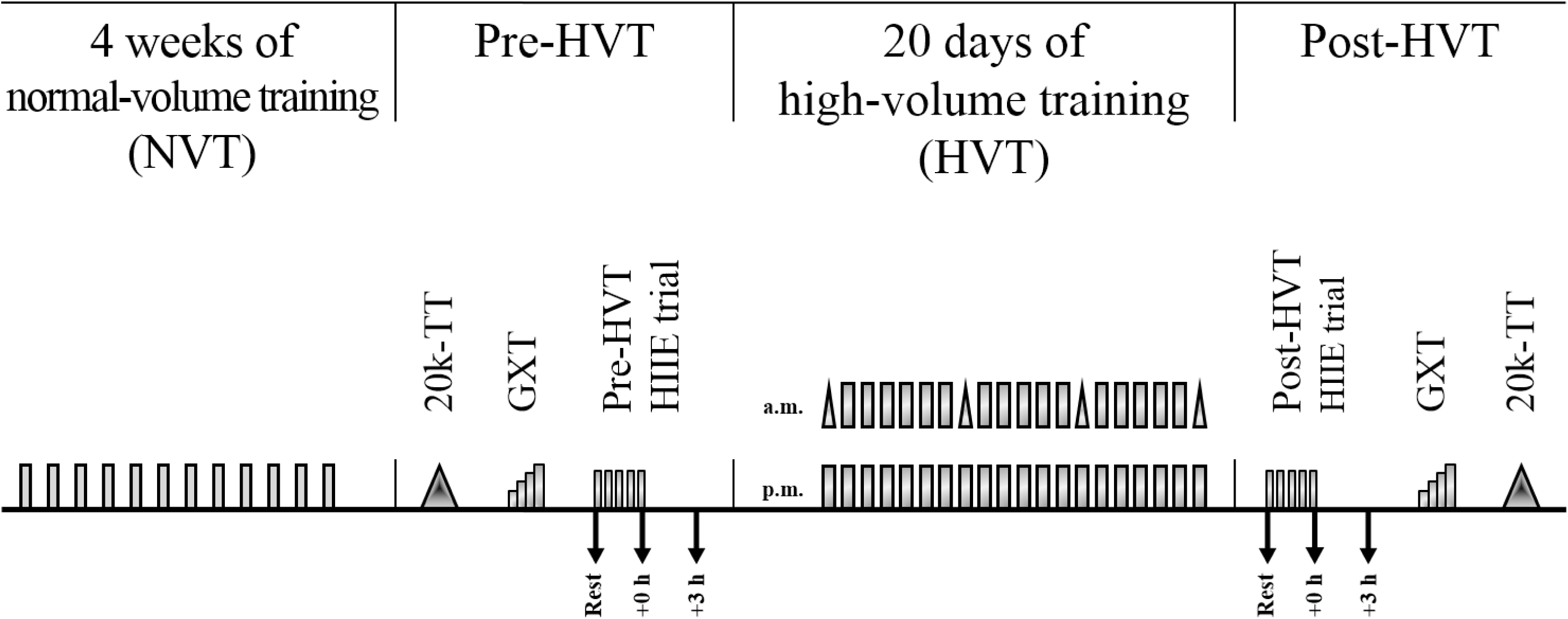
Study design. Grey rectangles indicate a HIIE session; grey triangles within HVT indicate a 10-km cycling time trial; each rectangle and/or vertical pair of rectangles and/or vertical pair of rectangles and triangles represents a training day; arrows indicate a skeletal muscle biopsy. Each test in both the Pre- and Post-HVT phase was separated by 48 to 72 hours. 20k-TT: 20-km cycling time trial; GXT: graded exercise test; HIIE: high-intensity interval exercise; Rest: skeletal muscle biopsy at rest; +0 h: skeletal muscle biopsy taken at the end of the HIIE session; +3 h: skeletal muscle biopsy taken three hours after the completion of the HIIE session.

#### 20k-TT

Cycling time trials were performed on an electronically-braked cycle ergometer (Velotron, RacerMate, USA) after a 6-min warm-up were participants cycled for 4 min at 66% of the power attained at the lactate threshold (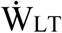), followed by 2 min at 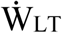, and 2 min of rest. During these tests, participants were only allowed access to cadence and completed distance. Heart rate was monitored (Polar-Electro, Finland) during all exercise trials and training sessions.

#### GXT

A discontinuous graded exercise test was performed on an electronically-braked cycle ergometer (Lode Excalibur, v2.0, The Netherlands) to determine peak oxygen uptake 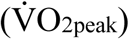, peak power 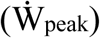, and 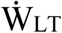(using the modified D_max_ method (5)), and the exercise intensity for both the biopsy trial and the HVT training sessions, as previously described (25). Briefly, the test began at 60, 90, or 120 W, depending on participants’ fitness levels, and was increased by 30 W every 4 min. Stages were interspersed with 30-s breaks for the measurement of fingertip capillary blood lactate concentration using a pre-calibrated blood-lactate analyzer (YSI 2300 STAT Plus, YSI, USA). Participants were instructed to keep a cadence above 60 rpm and were only allowed access to cadence and elapsed time; the GXT was terminated when participants reached volitional exhaustion or cadence dropped below 60 rpm. The 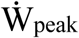 was determined as the power of the last completed stage plus 7.5 W for every additional minute completed. O_2_ and CO_2_ concentrations were analyzed from expired air using a pre-calibrated gas analyzer (Moxus 2010, AEI technologies, USA), and 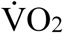 values were recorded every 15 s. The average of the two highest consecutive 15-s values was recorded as a participant’s 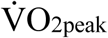. The same GXT was performed after 20 days of training to determine the relative exercise intensity of the Post-HVT biopsy trial.

#### Pre- and Post-HVT HIIE biopsy trials

Each participant performed the two biopsy trials in the morning and at the same time, to avoid variations caused by circadian rhythms. Participants were provided with a standardized dinner (55 kJ·kg^-1^ body mass (BM), providing 2.1 g carbohydrate·kg^-1^ BM, 0.3 g fat·kg^-1^ BM, and 0.6 g protein·kg^-1^ BM) and breakfast (41 kJ·kg^-1^ BM, providing 1.8 g carbohydrate·kg^-1^ BM, 0.2 g fat·kg^-1^ BM, and 0.3 g protein·kg^-1^ BM) to minimize variability in muscle gene and protein expression attributable to diet, as previously described (23). While participants rested in the supine position, and after injection of local anesthetic (1% xylocaine) into the skin and fascia of the vastus lateralis muscle, three small incisions were made about 2-3 cm apart. A resting muscle biopsy was taken (Rest) using a biopsy needle with suction. Approximately ten minutes later participants were helped to an electronically-braked cycle ergometer (Velotron, RacerMate, USA) and began a warm up consisting of cycling for four minutes at 66% of 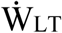, followed by 2 min at 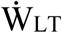, and 2 min of rest, after which the Pre-HVT HIIE session began. HIIE consisted of five 4-min intervals at an exercise intensity equal to 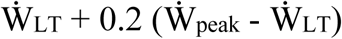, interspersed with two minutes of recovery at 60 W. Immediately after termination of HIIE (∼5 to 10 s), a second skeletal muscle biopsy was taken (+0 h), while a third one was obtained after three hours of recovery (+3 h), during which time participants were allowed access to water *ab libitum* and had no access to food. Skeletal muscle samples were rapidly cleaned of excess blood, fat, and connective tissue, were snap frozen in liquid nitrogen, and later stored at −80°C for subsequent analyses. By design, the Post-HVT HIIE biopsy trial was performed at the same absolute exercise intensity used during the Pre-HVT trial, and followed an identical format.

#### HVT

The day following the Pre-HVT HIIE biopsy trial participants began HIIT twice a day for 20 consecutive days. Training sessions were performed in the morning and afternoon and consisted of either 4- or 2-min intervals, interspersed with a 2- or 1-min recovery period at 60 W, respectively. To avoid stagnation, the training stimulus was progressively increased daily by virtue of increasing either the relative exercise intensity (from 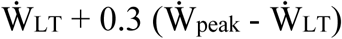 to 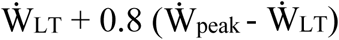 for the 4-min intervals, and from 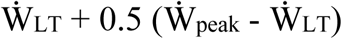 to 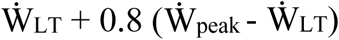 for the 2-min intervals), or the number of repetitions (from five to twelve bouts for the 4-min intervals, and from eight to twenty-two bouts for the 2-min intervals) (23). As a result, single-session duration increased from 30–35 min to 70–80 min. All participants progressively increased their relative exercise intensity and number of repetitions. A 10-km cycling time trial was performed before, and at regular weekly intervals during, the HVT to monitor participants for signs of overreaching, as previously described (23). The intention was to prevent overreaching by reducing the training load if performance decreased by more than 10% (28). However, no participants experienced a performance loss throughout the entire study, and the training protocol was completed as planned. All participants completed a minimum of 36 (equivalent to 90%) training sessions; average compliance was 96.5% of the prescribed number of sessions.

### Skeletal muscle analyses

#### Subcellular fractionation

Nuclear and cytosolic fractions were prepared from 35 to 50 mg of skeletal muscle using a commercially-available nuclear extraction kit (NE-PER, Pierce, USA). Briefly, muscle samples were washed in phosphate-buffered saline (PBS), homogenized in CER-I buffer containing a protease/phosphatase inhibitor cocktail (Cell Signaling Technology [CST], 5872) and centrifuged at ∼16,000 g. The supernatant was taken as the crude cytosolic fraction. The pellet containing nuclei was washed six times in PBS to minimize cytosolic contamination and nuclear protein was extracted by centrifugation (∼16,000 g) in a high-salt NER buffer supplemented with the same inhibitors cocktail and following the manufacturers’ instructions. Protein concentration was determined in triplicate using a commercial colorimetric assay (Bio-Rad Protein Assay kit-II; Bio-Rad, Gladesville, NSW, Australia). Nuclear and cytosolic fraction enrichment was confirmed by blotting the separated fractions against a nuclear (histone H3) and a cytosolic (lactate dehydrogenase A [LDHA]) protein; histone H3 was mainly detected in nuclear fractions, whereas LDHA was mainly detected in cytosolic fractions (Figure 2A), indicating the subcellular fractionation enrichment was successful.

**Figure 2.**
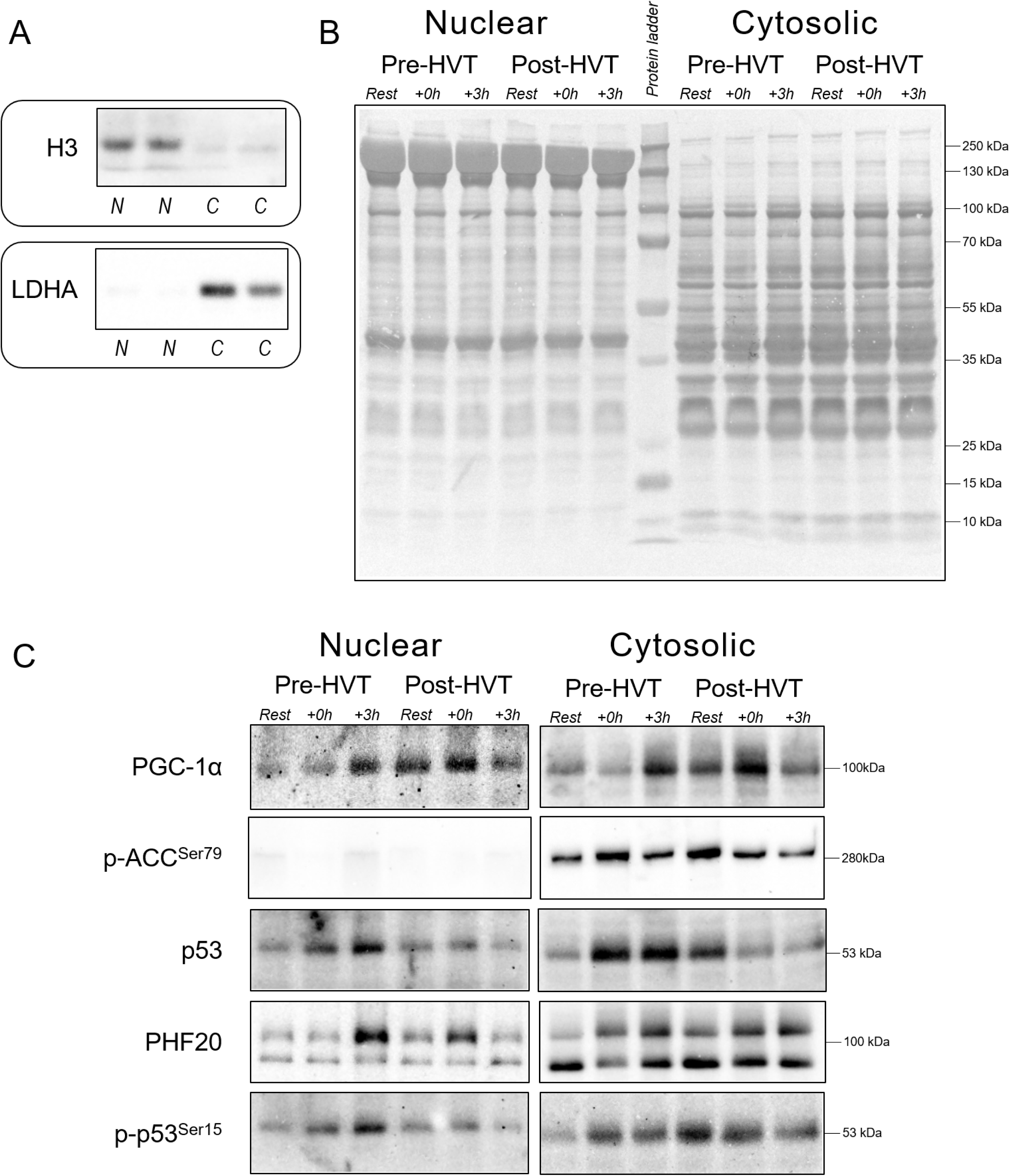
Representative immunoblots, subcellular enrichment and protein loading controls. (a) Representative immunoblots of peroxisome proliferator-activated receptor γ coactivator-1α (PGC-1α), acetyl-CoA carboxylase phosphorylated at serine 79 (p-ACC^Ser79^), p53, plant homeodomain finger-containing protein 20 (PHF20), and p53 phosphorylated at serine 15 (p-p53^Ser15^) measured in the nuclear and cytosolic fractions obtained from human vastus lateralis muscle biopsies, before (Rest), immediately post (+0 h), and 3 h (+3 h) after a single session of high-intensity interval exercise (HIIE) performed at the same absolute intensity before (Pre-HVT) and after (Post-HVT) 40 sessions of twice-daily high-volume high-intensity interval training (HVT). PHF20: top band at ∼105 kDa. No band was detected in the nuclear fractions for p-ACC^Ser79^. (b) Histone H3 and lactate dehydrogenase A (LDHA) were used as indicators of cytosolic and nuclear enrichment, respectively. N: nuclear fractions; C: cytosolic fractions. (c) Whole-lane Coomassie blue staining for both nuclear and cytosolic fractions was used to verify equal loading between lanes. The immunoblot and whole-lane Coomassie images in this figure were cropped to improve the conciseness and clarity of the presentation.

#### Immunoblotting

Muscle lysates (10 to 50 μg) were separated by electrophoresis using SDS-PAGE gels (8-15%) as previously described (24). An internal standard was loaded in each gel, and each lane was normalized to this value to reduce gel-to-gel variability. Whole-lane Coomassie blue staining (71) was performed to verify correct loading and equal transfer between lanes (Figure 2B). The following primary antibodies were used (supplier, catalogue number): histone H3 (CST, 9715), LDHA (CST, 2012), p53 (CST, 2527), p-acetyl-CoA carboxylase (p-ACC^Ser79^; CST, 3661), PGC-1α (Calbiochem, st-1202), plant homeodomain finger-containing protein 20 (PHF20; CST, 3934), and p-p53^Ser15^ (CST, 9284). Representative images for all target proteins are presented in Figure 2C.

#### Total RNA isolation

Total RNA was isolated from ∼15 mg of muscle tissue as previously described (14). Briefly, samples were homogenized (FastPrep FP120 Homogenizer; Thermo Savant) in the presence of 1 g of zirconia/silica beads (1.0 mm; Daintree Scientific, St. Helens, TAS, Australia) and 800 μL of TRIzol® Reagent (Invitrogen, Melbourne, Australia). Lysates were centrifuged at 13,000 rpm for 15 min at 4°C; the supernatant was collected, combined with chloroform (Sigma-Aldrich, St Louis, USA), and total RNA was extracted using the TRIzol® protocol as per manufacturer’s instructions. RNA precipitation was performed for at least 2 h at −20°C in the presence of 400 μL of isopropanol and 10 μL of 5 M NaCl (both Sigma-Aldrich, St Louis, USA). RNA concentration was determined spectrophotometrically (Nanodrop ND1000, Thermo Fisher Scientific, USA) by measuring the absorbance at 260 (A260) and 280 (A280) nm, with A260/A280 ratios > 1.8 indicating high-quality RNA (41). To ensure RNA was free of DNA contamination samples were DNase treated using an RQ1 RNase-free DNase kit (Promega Corporations, Madison, USA).

#### Real-time PCR (qPCR)

First-strand cDNA synthesis was performed on 300 ng of total RNA using a thermal cycler (S1000 Thermal Cycler; Bio-Rad; Bio-Rad, Gladesville, NSW, Australia) and the commercially available iScript™ cDNA synthesis kit (Bio-Rad, Gladesville, NSW, Australia) in the presence of random hexamers and oligo(dT)s, according to the manufacturer’s directions. Forward and reverse primers for all genes investigated (Table 1) were designed based on NCBI RefSeq using NCBI Primer-BLAST (www.ncbi.nlm.nih.gov/BLAST/), and specificity of the amplified product was confirmed by melting point dissociation curves. The mRNA expression of cytochrome *c* (cyt *c*), heat shock 70 kDa protein 1A (HSPA1A, usually referred to as HSP70), histone acetyltransferase KAT2A (KAT2A, usually referred to as general control of amino-acid synthesis 5 [GCN5]), nuclear respiratory factor 1 (NRF-1) and 2 (NRF-2), p53, PGC-1α, PHF20, peroxisome proliferator-activated receptor alpha (PPARα), peroxisome proliferator-activated receptor delta (PPARδ), peroxisome proliferator-activated receptor gamma (PPARγ), NAD-dependent protein deacetylase sirtuin-1 (SIRT1), and mitochondrial transcription factor A (TFAM) were quantified by quantitative real-time PCR (Mastercycler® RealPlex2, Eppendorf, Germany), using SYBR Green chemistry (iTaqTM Universal SYBR® Green Supermix; Bio-Rad, Gladesville, NSW, Australia) (10 µL PCR reaction volume). All samples were run in duplicate simultaneously with template free controls, using an automated pipetting system (epMotion 5070, Eppendorf, Germany) to reduce technical variation (41). The following PCR cycling patterns were used: initial denaturation at 95°C (3 min), 40 cycles of 95°C (15 s) and 60°C (60 s). Relative changes in mRNA content were calculated using the 2^−ΔΔCt^ method. To account for the efficiency of RT and initial RNA concentration, the mRNA expression of four housekeeping genes was quantified, and their stability was determined using the BestKeeper software (56). Cyclophilin, glyceraldehyde 3-phosphate dehydrogenase (GAPDH), and beta-2-microglobulin (B2M) were classified as stable, whereas TATA-binging protein (TBP) was reported as unstable and was therefore excluded. These results were confirmed by the Normfinder algorithm (2).

**Table 1.**
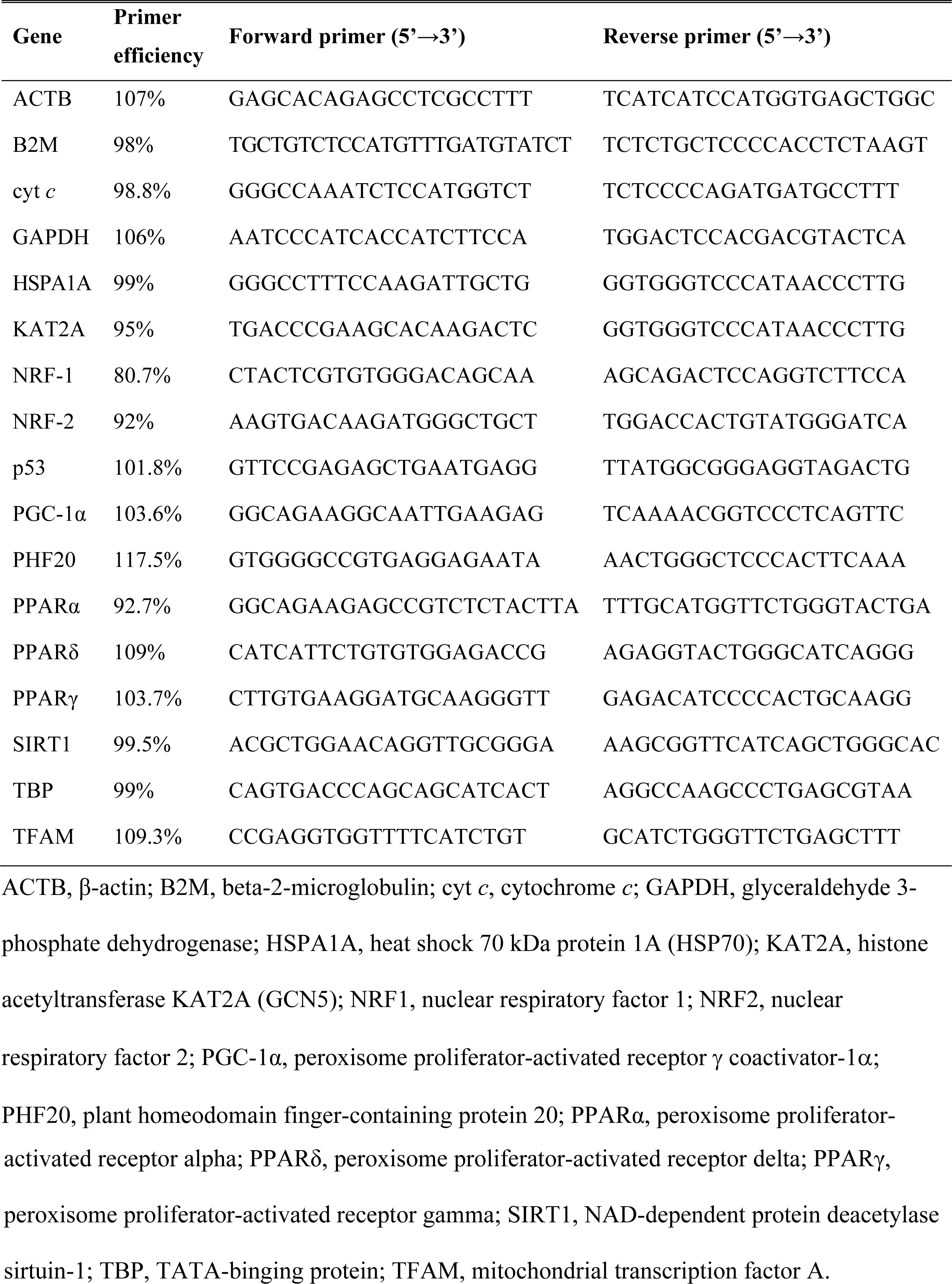
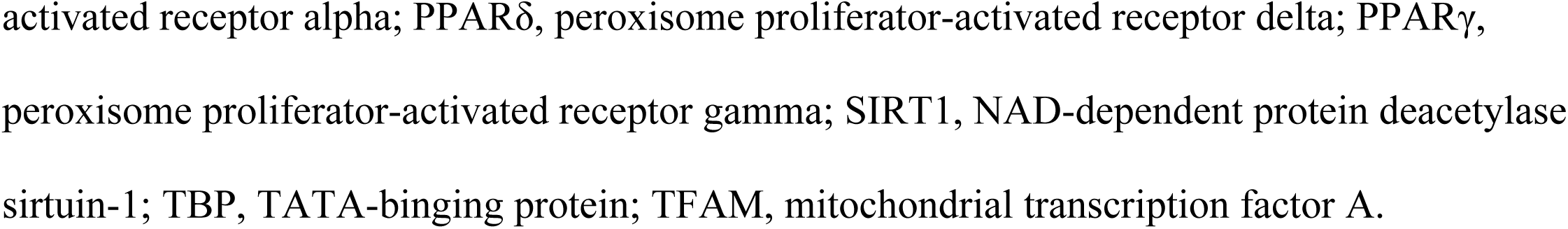
Primers used for real-time PCR analyses of mRNA expression.

#### Statistical analysis

All values are reported as mean ± SD unless otherwise specified. Outliers (defined as values outside the mean ± 3SD) were first removed. Normality was assessed with a Shapiro-Wilk test; datasets that failed the normality test (*P* < 0.05) were log transformed, and if the dataset was still non-normal the reciprocal value was used. To investigate the influence of exercise (Rest, +0 h, and +3 h) and training (Pre-HVT and Post-HVT), and the interaction between these two variables, two-way ANOVA with repeated measures were performed. Interactions were followed by Tukey’s honestly significant difference post-hoc tests to assess differences between time points (both within and between trials). In addition, main effects of exercise were further analyzed with pre-planned contrasts comparing the effect of exercise within biopsy trials only. Resting protein and mRNA content values in the Pre- and Post-HVT trials were also compared with a pre-planned paired t-test, to determine the effects of 40 sessions of HIIT. SigmaPlot 13.0 software (Jandel Scientific, USA) was used for all statistical analyses. The level of statistical significance was set a priori at *P* < 0.05.

## Results

### Total work during the biopsy trials

By design, the Pre- and Post-HVT HIIE sessions were performed at the same absolute exercise intensity (231.1 ± 33.1 W, Figure 3) and resulted in the same total work (277.3 ± 39.8 kJ). There was an increase (9.0 ± 6.1%, *P* = 0.002) in the power attained at the lactate threshold (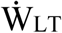) following training (215.5 ± 32.2 *vs.* 234.7 ± 36.8 W, Pre- and Post-HVT, respectively; Figure 3), which resulted in the relative exercise intensity of the Pre-HVT biopsy trial (107.4 ± 1.2% of 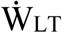) being greater than the Post-HVT biopsy trial (98.8 ± 5.2% of 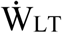). Following training, there was also an increase in peak power 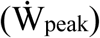 (7.8 ± 4.4%, *P* = 0.001; 292.5 ± 37.9 *vs.* 315.2 ± 42.3 W, Pre- and Post-HVT, respectively; Figure 3); consequently, the exercise intensity expressed relative to 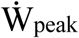 was also greater in the Pre-HVT biopsy trial (78.9 ± 2.4% of 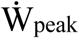) than the Post-HVT biopsy trial (73.3 ± 3.7% of 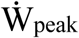). Post-HVT, there was an increase in peak oxygen uptake 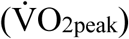 (11.7 ± 7.6%, *P* = 0.001; 46.2 ± 7.6 *vs.* 51.4 ± 7.8 mL · min^-1^ · kg^-1^, Pre- and Post-HVT, respectively), whereas 20-km cycling time trial (20k-TT) time was decreased (5.2 ± 2.3%, *P* < 0.001; 2140.8 ± 99.9 *vs.* 2028.1 ± 87.5 s, Pre- and Post-HVT, respectively). The participants’ BM did not change post training (0.2 ± 1.6 %, *P* = 0.720; 80.4 ± 14.8 *vs.* 80.6 ± 14.5 kg, Pre- and Post-HVT, respectively).

**Figure 3.**
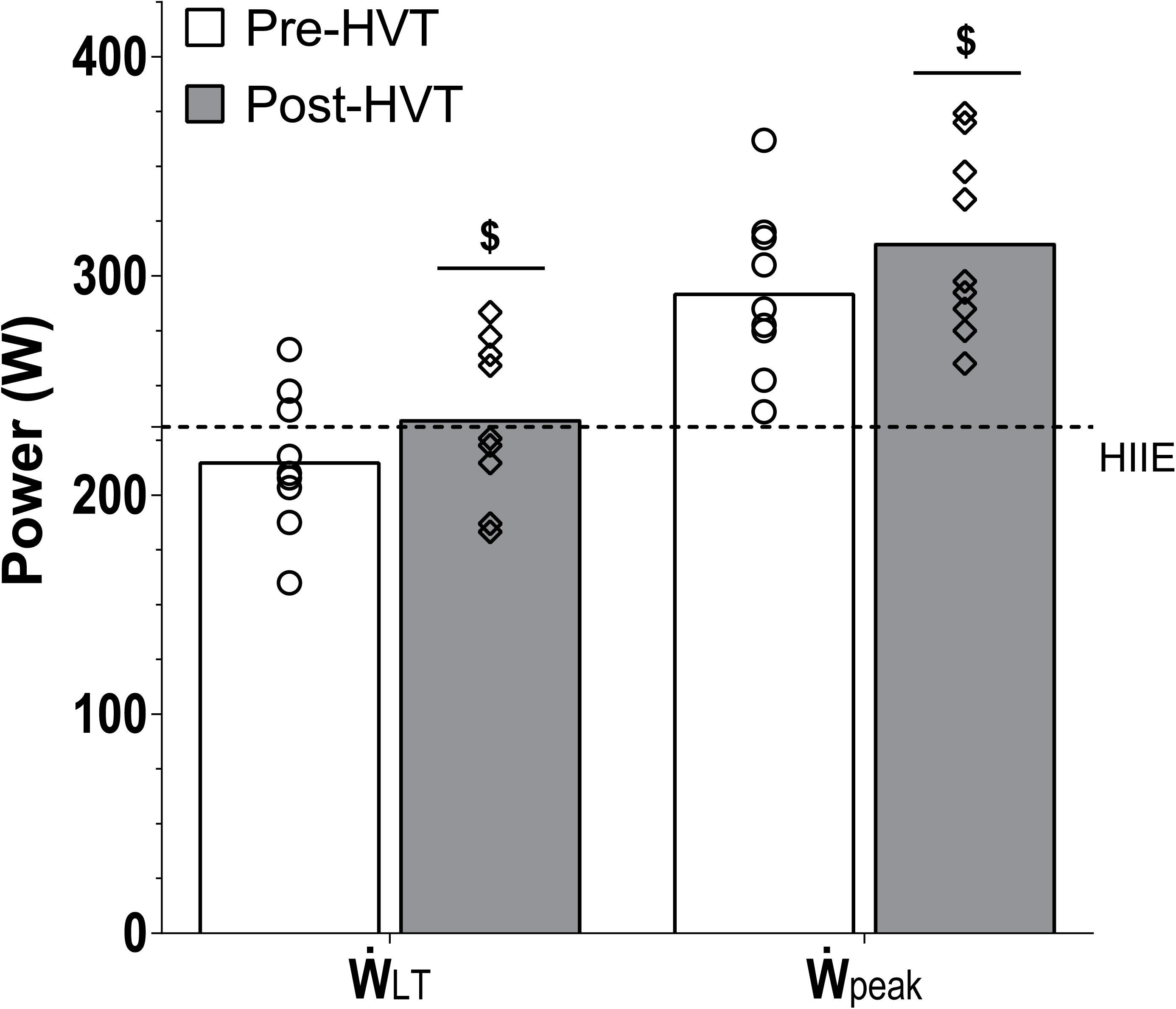
Power attained at the lactate threshold 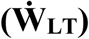, peak power achieved during the graded exercise test 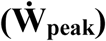, and mean power of the Pre- and Post-HVT high-intensity interval exercise (HIIE) biopsy trials. 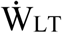 and 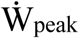 were assessed before (Pre-HVT) and after (Post-HVT) 40 sessions of twice-daily high-volume high-intensity interval training (HVT). Open circles (Pre-HVT) and open diamonds (Post-HVT) represent individual values; white (Pre-HVT) and grey (Post-HVT) bars represent mean values; the dotted line represents the mean power during the Pre- and Post-HVT HIIE biopsy trials. n = 9. ^$^ *P* < 0.05 *vs.* Pre-HVT Rest by paired t-test.

### Muscle analyses

Representative immunoblots are presented in Figure 2C.

#### PGC-1α protein content

There was an interaction effect in both subcellular compartments (nucleus: *P* = 0.044, cytosol: *P* = 0.004). In the nucleus (Figure 4A), PGC-1α was increased at +3 h compared with Rest during the Pre-HVT (3.1-fold, *P* = 0.002), but not during the Post-HVT (1.0-fold, *P* = 0.869) biopsy trial. During Pre-HVT, nuclear PGC-1α was also greater at +3 h compared with Post-HVT (3.1-fold, *P* = 0.015). At Rest, nuclear PGC-1α protein content was greater Post-HVT compared with Pre-HVT (1.8-fold, *P* = 0.013).

**Figure 4.**
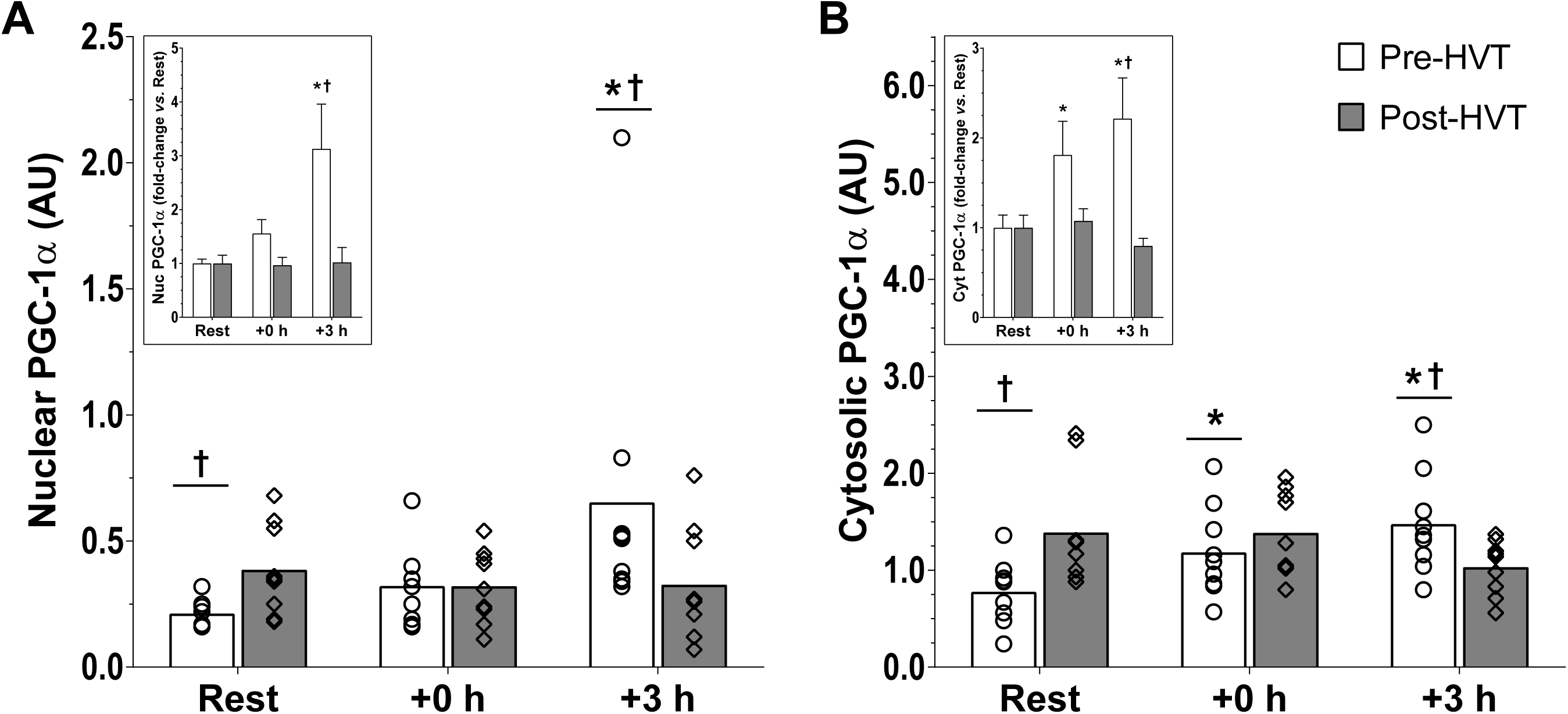
Peroxisome proliferator-activated receptor γ coactivator-1α (PGC-1α) protein. Protein content of PGC-1α in nuclear (a), and cytosolic (b) sub fractions before (Rest), immediately post (+0 h), and 3 h (+3 h) after a single session of high-intensity interval exercise (HIIE) performed at the same absolute intensity before (Pre-HVT) and (Post-HVT) 40 sessions of twice-daily high-volume high-intensity interval training (HVT), in the vastus lateralis muscle of young healthy men (n = 9). Open circles (Pre-HVT) and open diamonds (Post-HVT) represent individual values; white (Pre-HVT) and grey (Post-HVT) bars represent mean values. * *P* < 0.05 *vs.* Rest of the same group; ^†^ *P* < 0.05 *vs.* same time point of Post-HVT trial by two-way ANOVA with repeated measures followed by Tukey’s honestly significant difference post-hoc test, or pre-planned paired t-test for Rest values between trials. To more clearly depict fold-changes in post-exercise values from potentially different Rest values in the untrained and trained state, an inset has been added to each main figure (note, significant differences between trained and untrained values at Rest are not reported in insets as these values are both normalized to 1); the error bars represent the standard error of the mean (SEM).

In the cytosol (Figure 4B), PGC-1α increased compared with Rest both at +0 h (1.8-fold, *P* = 0.036) and +3 h (2.2-fold, *P* < 0.001) during the Pre-HVT, but not during the Post-HVT (1.1-fold, *P* = 1.000 at +0 h; 0.8-fold, *P* = 0.070 at +3 h) biopsy trial. During the Pre-HVT biopsy trial, cytosolic PGC-1α was also greater at +3 h (1.5-fold, *P* = 0.017) compared with the same time point of the Post-HVT biopsy trial. At Rest, cytosolic PGC-1α was greater Post-HVT compared with Pre-HVT (2.0-fold, *P* = 0.005).

#### Gene expression

There was an interaction effect for PGC-1α mRNA content (*P* = 0.020; Figure 5A), which was increased at +3 h compared with Rest during the Pre-HVT (3.6-fold, *P* < 0.001), but not during the Post-HVT (2.0-fold, *P* = 0.129) biopsy trial. During the Pre-HVT biopsy trial, the mRNA content of PGC-1α at +3 h was also greater (1.9-fold, *P* < 0.001) compared with that recorded at the same time point during the Post-HVT biopsy trial. There was no change in p53 mRNA content throughout (interaction: *P* = 0.425; main effect of exercise: *P* = 0.379; Figure 5B). Results for the mRNA content of cyt *c*, GCN5, HSP70, NRF-1 and NRF-2, PHF20, PPARα, PPARδ, PPARγ, SIRT1, and TFAM are reported in Table 2.

**Table 2.**
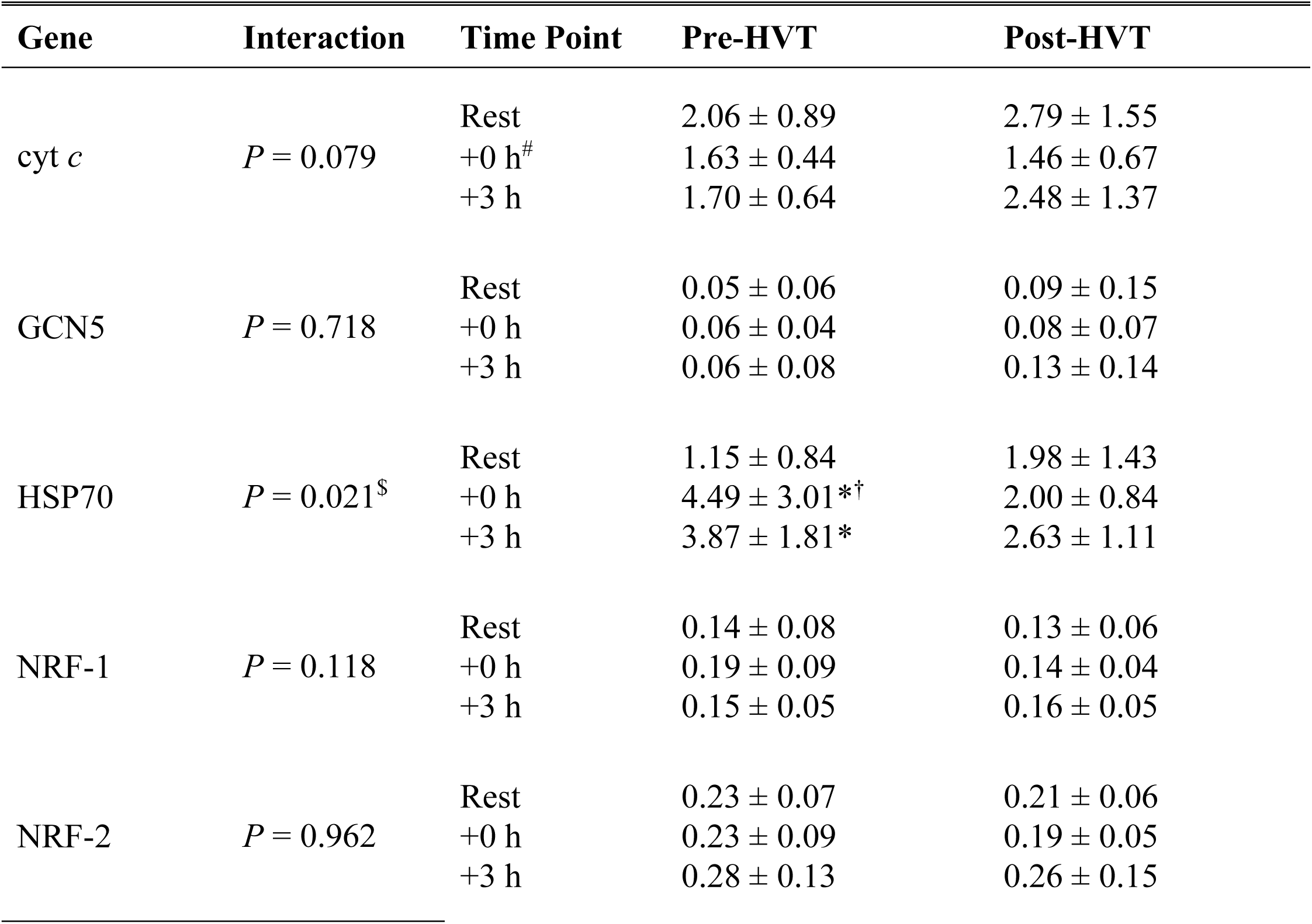
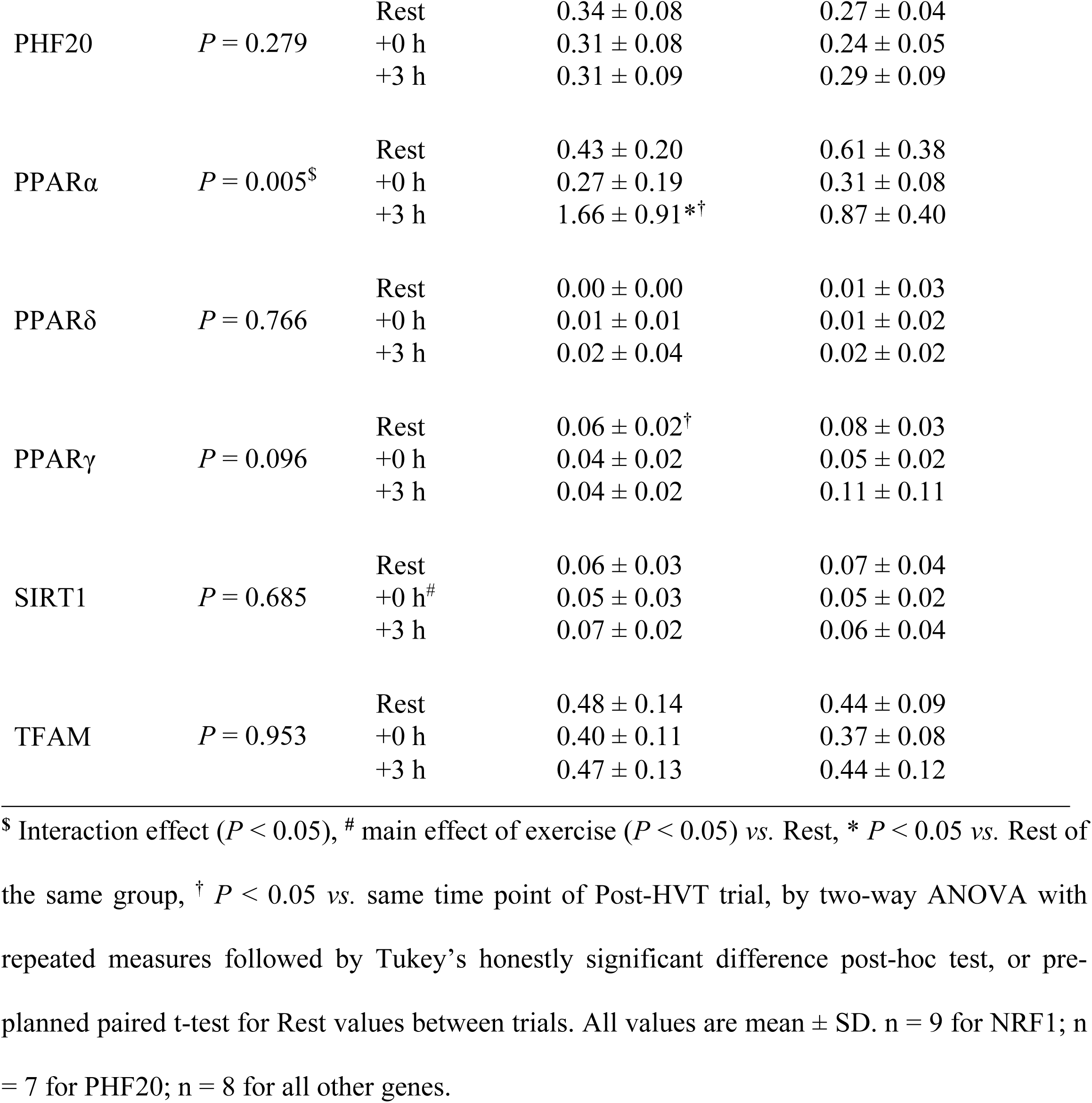
mRNA content measured in whole tissue of cytochrome *c* (cyt *c*), histone acetyltransferase KAT2A (GCN5), heat shock 70 kDa protein 1A (HSP70), nuclear respiratory factor 1 (NRF-1) and 2 (NRF-2), plant homeodomain finger-containing protein 20 (PHF20), peroxisome proliferator-activated receptor alpha (PPARα), peroxisome proliferator-activated receptor delta (PPARδ), peroxisome proliferator-activated receptor gamma (PPARγ), NAD-dependent protein deacetylase sirtuin-1 (SIRT1), and mitochondrial transcription factor A (TFAM) measured immediately post (+0 h) and 3 h (+3 h) after a single session of high-intensity interval exercise (HIIE) performed at the same absolute intensity before (Pre-HVT) and after (Post-HVT) 40 sessions of twice-daily high-volume high-intensity interval training (HVT), in the vastus lateralis muscle of young healthy men. Values are expressed relative to TATA-binging protein (TBP), glyceraldehyde 3-phosphate dehydrogenase (GAPDH), and β-actin (ACTB) housekeeping genes.

**Figure 5.**
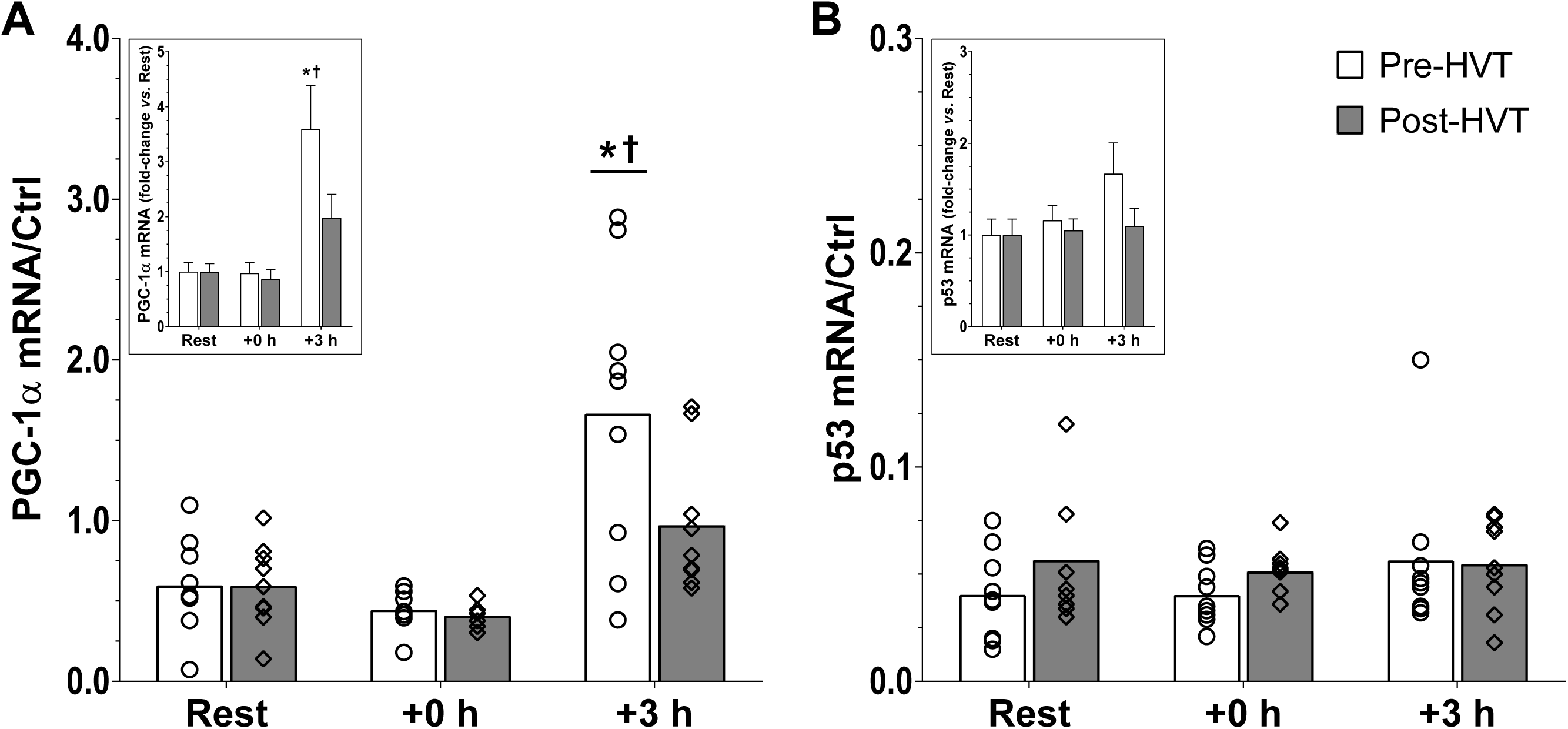
Gene expression in whole tissue. mRNA content of peroxisome proliferator-activated receptor γ coactivator-1α (PGC-1α) (a), and p53 (b) before (Rest), immediately post (+0 h), and 3 h (+3 h) after a single session of high-intensity interval exercise (HIIE) performed at the same absolute intensity before (Pre-HVT) and after (Post-HVT) 40 sessions of twice-daily high-volume high-intensity interval training (HVT), in the vastus lateralis muscle of young healthy men (n = 9). Values are expressed relative to TATA-binging protein (TBP), glyceraldehyde 3-phosphate dehydrogenase (GAPDH), and β-actin (ACTB) housekeeping genes (Ctrl in the figure). Open circles (Pre-HVT) and open diamonds (Post-HVT) represent individual values; white (Pre-HVT) and grey (Post-HVT) bars represent mean values. * *P* < 0.05 *vs.* Rest of the same group; ^†^ *P* < 0.05 *vs.* same time point of Post-HVT trial by two-way ANOVA with repeated measures followed by Tukey’s honestly significant difference post-hoc test, or by pre-planned paired t-test for Rest values between trials. To more clearly depict fold-changes in post-exercise values from potentially different Rest values in the untrained and trained state, an inset has been added to each main figure; the error bars represent the SEM.

#### Phosphorylation of acetyl-CoA carboxylase (ACC) at serine 79 (p-ACC^Ser79^) protein content

p-ACC^Ser79^ was not detected in nuclear fractions (Figure 2C). In the cytosol (Figure 6), no interaction effect was reported (*P* = 0.774); however, there was a main effect of exercise (*P* < 0.001), whereby p-ACC^Ser79^ was greater compared with Rest at +0 h (1.7-fold, *P* < 0.001). Pre-planned comparisons within biopsy trials indicated that at +0 h cytosolic p-ACC^Ser79^ was greater compared with Rest during the Pre-HVT (2.0-fold, *P* = 0.013), but not during the Post-HVT (1.4-fold, *P* = 0.114) biopsy trial.

**Figure 6.**
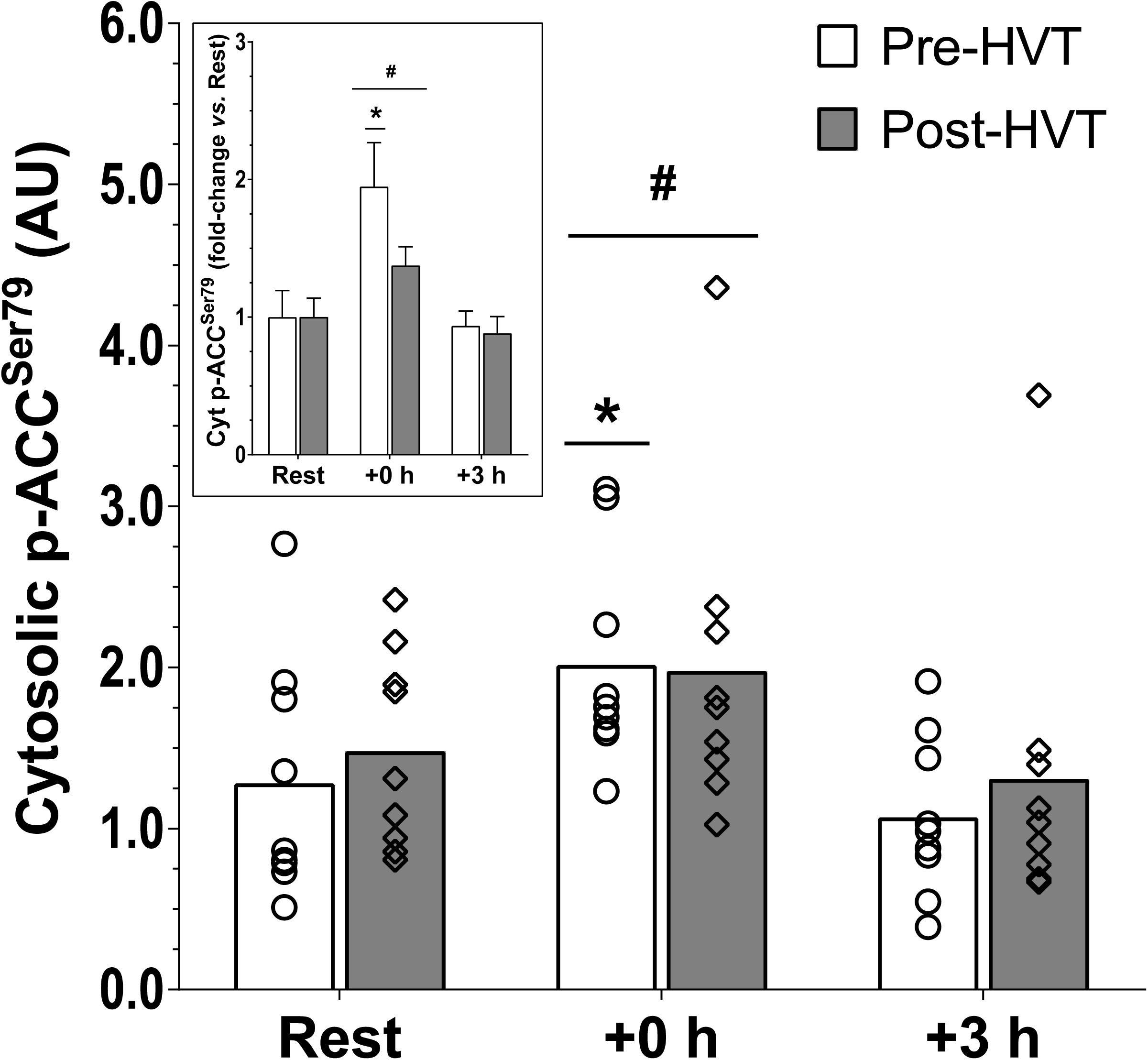
Phosphorylation of acetyl-CoA carboxylase (ACC) at serine 79 (p-ACC^Ser79^). Protein content of cytosolic p-ACC^Ser79^ before (Rest), immediately post (+0 h), and 3 h (+3 h) after a single session of high-intensity interval exercise (HIIE) performed at the same absolute intensity before (Pre-HVT) and after (Post-HVT) 40 sessions of twice-daily high-volume high-intensity interval training (HVT), in the vastus lateralis muscle of young healthy men (n = 9). Open circles (Pre-HVT) and open diamonds (Post-HVT) represent individual values; white (Pre-HVT) and grey (Post-HVT) bars represent mean values. ^**#**^ main effect of exercise (*P* < 0.05) *vs.* Rest; * *P* < 0.05 vs. Rest of the same group by two-way ANOVA with repeated measures followed by Tukey’s honestly significant difference post-hoc test, or by pre-planned paired t-test for Rest values between trials. To more clearly depict fold-changes in post-exercise values from potentially different Rest values in the untrained and trained state, an inset has been added to each main figure; the error bars represent the SEM.

#### p53 protein content

In the nucleus (Figure 7A), there was an interaction effect (*P* = 0.016); nuclear p53 was increased at +3 h compared with Rest during the Pre-HVT (2.8-fold; *P* = 0.004), but not during the Post-HVT (1.2-fold, *P* = 0.328) biopsy trial. At Rest, nuclear p53 was greater Post-HVT compared with Pre-HVT (1.6-fold, *P* = 0.038).

**Figure 7.**
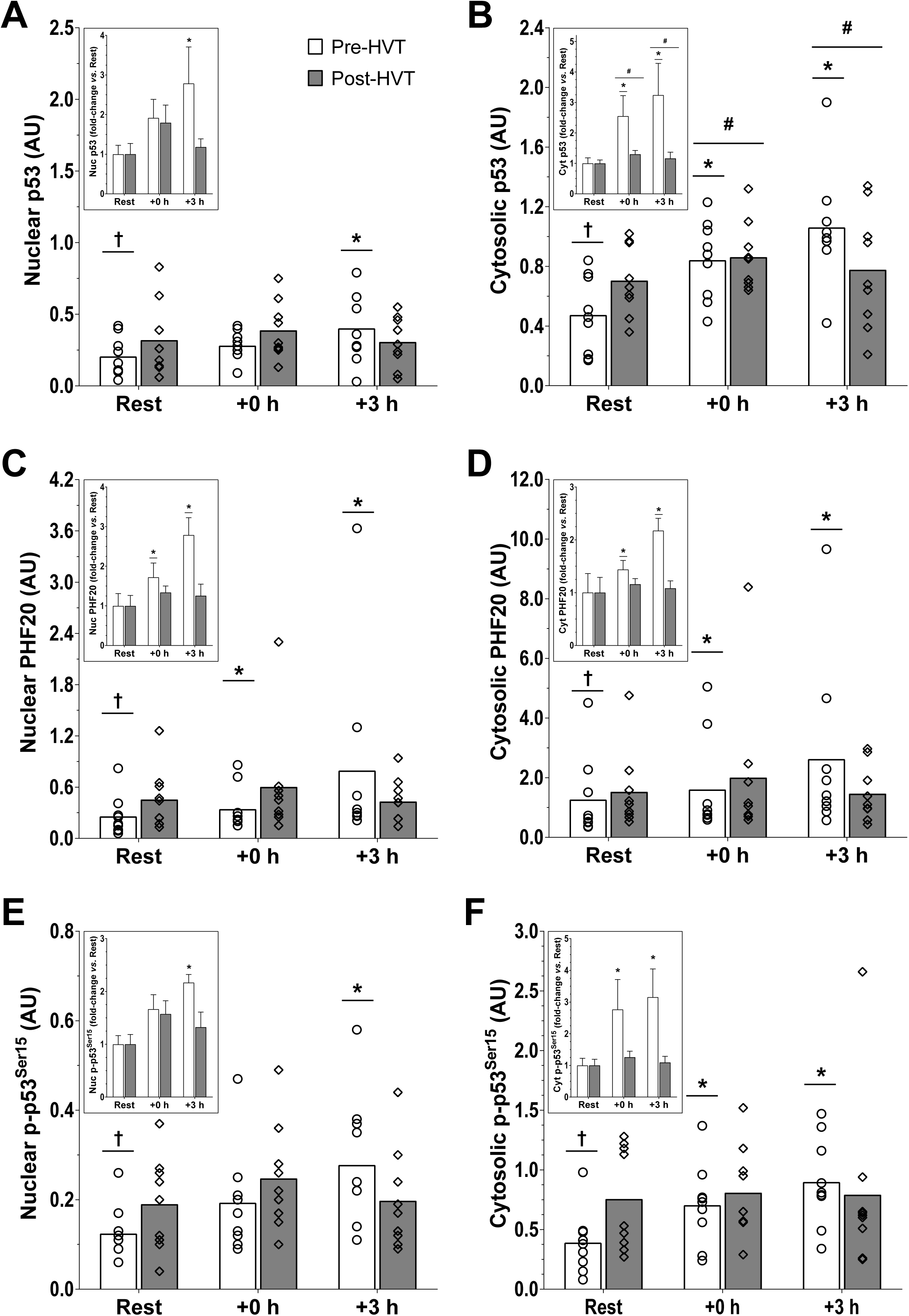
p53 and plant homeodomain finger-containing protein 20 (PHF20) protein. Protein content of nuclear (a) and cytosolic (b) p53, of nuclear (c) and cytosolic (d) PHF20, and of nuclear (e) and cytosolic (f) p-p53^Ser15^ assessed before (Rest), immediately post (+0 h), and 3 h (+3 h) after a single session of high-intensity interval exercise (HIIE) performed at the same absolute intensity before (Pre-HVT) and after (Post-HVT) 40 sessions of twice-daily high-volume high-intensity interval training (HVT), in the vastus lateralis muscle of young healthy men (n = 9). Open circles (Pre-HVT) and open diamonds (Post-HVT) represent individual values; white (Pre-HVT) and grey (Post-HVT) bars represent mean values. ^**#**^ main effect of exercise (*P* < 0.05) *vs.* Rest; * *P* < 0.05 *vs.* Rest of the same group; ^†^ *P* < 0.05 *vs.* same time point of Post-HVT trial by two-way ANOVA with repeated measures followed by Tukey’s honestly significant difference post-hoc test, or by pre-planned paired t-test for Rest values between trials. To more clearly depict fold-changes in post-exercise values from potentially different Rest values in the untrained and trained state, an inset has been added to each main figure (note, significant differences between trained and untrained values at Rest are not reported in insets as these values are both normalized to 1); the error bars represent the SEM.

In the cytosol (Figure 7B), the interaction effect was not statistically significant (*P* = 0.051); however, there was a main effect of exercise (*P* = 0.003). Cytosolic p53 increased compared with Rest both at +0 h (1.9-fold, *P* = 0.019) and +3 h (2.2-fold, *P* = 0.004). Pre-planned comparisons within trials revealed that during the Pre-HVT biopsy trial cytosolic p53 was greater compared with Rest at both +0 h (2.6-fold, *P* = 0.020) and +3 h (3.2-fold, *P* < 0.001); however, during the Post-HVT biopsy trial no differences compared with Rest were reported at +0h (1.3-fold, *P* = 0.440) and +3 h (1.2-fold, *P* = 0.835). At Rest, cytosolic p53 was greater Post-HVT compared with Pre-HVT (1.9-fold, *P* = 0.015).

#### PHF20 protein content

There was an interaction effect in both subcellular compartments (nucleus: *P* = 0.019, cytosol: *P* = 0.025). In the nucleus (Figure 7C), PHF20 increased compared with Rest both at +0 h (1.7-fold, *P* = 0.016) and +3 h (2.8-fold, *P* < 0.001) during the Pre-HVT, but not during the Post-HVT (1.3-fold, *P* = 0.616 at +0 h; 1.3-fold, *P* = 0.858 at +3 h) biopsy trial. At Rest, nuclear PHF20 was greater Post-HVT compared with Pre-HVT (1.9-fold, *P* = 0.004).

In the cytosol (Figure 7D), PHF20 increased compared with Rest both at +0 h (1.4-fold, *P* = 0.032) and +3 h (2.2-fold, *P* < 0.001) during the Pre-HVT, but not during the Post-HVT (1.2-fold, *P* = 0.890 at +0 h; 1.1-fold, *P* = 0.996 at +3 h) biopsy trial. At Rest, cytosolic PHF20 was greater Post-HVT compared with Pre-HVT (1.5-fold, *P* = 0.013).

#### p-p53^Ser15^ protein content

In the nucleus (Figure 7E), there was an interaction effect (*P* = 0.021); nuclear p-p53^Ser15^ was increased compared with Rest at +3 h during the Pre-HVT (2.2-fold; *P* = 0.001), but not during the Post-HVT (1.3-fold, *P* = 0.970) biopsy trial. At Rest, nuclear p-p53^Ser15^ was greater Post-HVT compared with Pre-HVT (1.5-fold, *P* = 0.043)

An interaction effect (*P* = 0.033) was also reported in the cytosol (Figure 7F). During Pre-HVT, cytosolic p-p53^Ser15^ was greater compared with Rest at both +0 h (2.8-fold, *P* = 0.018) and +3 h (3.2-fold, *P* < 0.001), but not during the Post-HVT (1.3-fold, *P* = 0.847 and 1.1-fold, *P* = 0.997 at +0 and +3 h, respectively) biopsy trial. At Rest, cytosolic p-p53^Ser15^ was greater Post-HVT compared with Pre-HVT (2.4-fold, *P* = 0.008).

## Discussion

We report that 40 sessions of HIIT resulted in the loss of all measured exercise-induced molecular changes recorded Pre-HVT. Although training-induced blunting of specific exercise-induced molecular adaptations in whole cell lysates has previously been reported (48, 49, 55, 69), this is the first study demonstrating training-induced blunting of selected markers of mitochondrial biogenesis at the subcellular level. Despite exercise-induced increases in both the nuclear and cytosolic fractions in PGC-1α, p53, PHF20, and p-p53^Ser15^ protein content prior to the HVT, there were no significant changes in any of these parameters when a session of HIIE was repeated at the same absolute exercise intensity post training. However, post-HVT there was an increase in resting values of most proteins measured in this study. In contrast to our findings, where exercise-induced upregulation of all measured parameters was blunted post-training, previous research has reported that training-induced blunting of exercise-induced molecular changes is not universal (55, 69). These discrepancies may relate to the much greater number of training sessions in the present study (40 *vs.* 7 (55) and 10 (69), respectively).

We observed a significant exercise-induced increase in PGC-1α protein content in both the nuclear and cytosolic fractions Pre-HVT, consistent with most previous research (24, 33, 43, 44). However, for the first time we report that these exercise-induced increases were absent post training in both subcellular fractions. A possible explanation for our findings is that Post-HVT the relative exercise intensity elicited during the session was lower compared with Pre-HVT (98.8 *vs.* 107.4% of 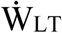 for Post- and Pre-HVT, respectively), suggesting that metabolic perturbations may also have been reduced post-training. This is supported by absence of significant changes in cytosolic p-ACC^Ser79^ Post-HVT (as described more in detail below) It has been proposed that metabolic perturbations (e.g., increases in intracellular calcium [Ca^2+^], adenosine monophosphate [AMP] to adenosine triphosphate [ATP] ratio, oxidized nicotinamide adenine dinucleotide [NAD^+^] to NADH ratio, reactive oxygen species [ROS] production) provide an important stimulus for exercise-induced mitochondrial biogenesis (17), and promote an increase in the nuclear content of PGC-1α protein (73).

The reported increase in PGC-1α protein content in both the nucleus and the cytosol during the Pre-HVT trial may be attributable, at least in part, to increased protein stability (63). Both p38 mitogen-activated protein kinase (MAPK) (61) and AMP-activated protein kinase (AMPK) (7) act as signaling proteins that increase PGC-1α stability via phosphorylation (and they act in similar fashion to also increase the stability of the p53 protein (39, 67)). Due to the limited amount of enriched lysates obtained during subcellular fractionation, we could not measure phosphorylation of p38 MAPK and/or AMPK directly. However, due to its molecular weight (∼280 kDa), when blotting for lower molecular weight proteins we were also able to measure p-ACC^Ser79^, a downstream target and commonly used marker of AMPK activation (9, 10, 38). As previously reported, p-ACC^Ser79^ was not detected in nuclear fractions (24, 44). However, we were able to measure cytosolic p-ACC^Ser79^ and make the novel observation that despite a post-exercise increase Pre-HVT, there was no significant change Post-HVT. This suggests that abrogation of AMPK signaling may have contributed to the abrogation of exercise-induced increases in PGC-1α (and p53) protein content post-training. Subcellular translocation is another factor that has been associated with increased PGC-1α protein content in the nucleus (73). While our data do not seem to indicate cytosolic/nuclear shuttling of PGC-1α, protein translocation is a complex series of cellular processes that cannot be assessed by subcellular fractionation coupled with the immunoblotting technique (1).

PGC-1α has been shown to be activated via deacetylation by SIRT1 (7, 63). Although previous research has reported exercise-induced increases in SIRT1 mRNA in human skeletal muscle following both low-intensity continuous (13) and high intensity interval (15) exercise, our results indicate a small significant decrease at +0 h (0.8-fold change). SIRT1 activity rather that protein content seems to regulate mitochondrial biogenesis in humans (27); however, due to limited tissue availability, we were not able to perform this measurement. It has also been reported that SIRT1 deacetylase activity may not be required for exercise-induced mitochondrial biogenesis or PGC-1α deacetylation, and that changes in the acetyltransferase activity and subcellular location of GCN5, a negative regulator of PGC-1α (18), may be more important factors regulating exercise-induced PGC-1α activity (57). Consistent with previous findings (15), we report no change in GCN5 mRNA content both Pre- and Post-HVT. Limited skeletal muscle availability precluded us from assessing GCN5 activity or protein content in different subcellular fractions.

The PGC-1α protein itself has been reported to stimulate PGC-1α transcriptional activity via an autoregulatory loop that requires coactivation of the myocyte enhancer factor-2 protein (30). The exercise-induced increase in PGC-1α mRNA content observed Pre-HVT is consistent with previous findings investigating HIIE (12, 15, 49, 51, 55, 59, 60) and with the notion that increased nuclear PGC-1α protein content and stability is associated with greater PGC-1α transcriptional activity (3). No exercise-induced increase in PGC-1α mRNA content was reported Post-HVT, suggesting that 20 days of HVT also blunted the exercise-induced increase in PGC-1α transcription. However, previous studies have reported a reduction (rather than complete loss) of the exercise-induced upregulation of PGC-1α mRNA content post-training compared to pre-training when the exercise session was repeated at the same relative (55) or absolute (49, 69) exercise intensity. This discrepancy may relate to the much greater number of sessions performed between exercise biopsy trials in our study compared with these three previous studies (40 *vs.* 7 to 12, respectively), and a likely greater reduction in the relative exercise intensity between the Pre- and Post-HVT trials. Moreover, in contrast to the three previous studies, our participants were habituated to HIIE; this raises the possibility that the greater molecular response recorded pre-training in the previous studies may be partly attributable to the “first bout” effect (4, 50).

To better characterize the effect of 40 sessions of HIIT on exercise-induced mitochondrial adaptations to HIIE, we measured the mRNA content of nuclear (NRF-1 and NRF-2 (65)) and mitochondrial (TFAM (66)) transcription factors regulating mitochondrial biogenesis that are transcriptionally controlled by PGC-1α (74). The mRNA content of cyt *c* (a gene under the regulation of PGC-1α and NRF1 (74)), p53 (a transcriptional regulator of PGC-1α gene expression (36)) and PHF20 (a transcription factor that activates p53 gene expression (54)), were also measured. In addition, we also assessed the mRNA content of three PPAR genes, which are involved in fatty acid metabolism and transport (19), and HSP70, a chaperone protein required for the import and folding of mitochondrial proteins (40). Both HSP70 and PPARα were increased following exercise during the Pre-HVT trial, but not during the Post-HVT trial, following a similar response to the majority of the molecular events linked with exercise-induced mitochondrial biogenesis measured in our study. Aside from a decrease in cyt *c* mRNA content at +0 h in both HIIE trials, we observed no exercise-induced changes in any of the other genes either Pre- or Post-HVT. It is important to note that a possible explanation for the lack of exercise-induced upregulation of some of these mRNAs (at least Pre-HVT) may relate to the biopsy timings chosen post-exercise, as there is evidence that the exercise-induced upregulation of some of these genes peaks more than 3 hours post-exercise (8, 12, 16, 21, 29, 54).

Similar to our results for PGC-1α protein, we observed an exercise-induced increase in p53 protein content pre-training in the nuclear and cytosolic fractions - as previously demonstrated (24, 70). However, this exercise-induced increase in both subcellular fractions was blunted following 40 training sessions. No other study has investigated exercise-induced changes in p53 protein content pre- and post-training in subcellular fractions. Nonetheless, our results are consistent with findings showing reduced/blunted exercise-induced mitochondrial adaptations (e.g., PGC-1α mRNA, PGC-1α protein in whole-muscle lysates) when the same exercise session is repeated post-training both at the same absolute (49, 69) or relative (55) exercise intensity.

A possible factor contributing to the lack of exercise-induced changes in nuclear p53 protein content Post-HVT is that 40 sessions of HIIT increased the resting values of p53 in in both fractions. A second factor relates to a possible decrease in subcellular shuttling (20); however, simply immunoblotting subcellular enriched fractions for p53 (or PGC-1α) protein is not a valid technique to demonstrate p53 (similar to PGC-1α) nuclear/cytosolic shuttling - a process requiring an intricate and tightly synchronized series of events (20, 47). Nonetheless, a further novel observation is that there was a concomitant increase in p53 and PHF20 protein content in both subcellular fractions Pre-HVT, but not Post-HVT. In this regard, PHF20 has been reported to increase p53 protein stability (53) by disrupting the murine double minute-2 (MDM2)-p53 interaction (11) responsible for p53 protein degradation (32, 53). Although we were not able to measure the interaction between these two proteins due to limited lysate availability, it is plausible that our findings may indicate greater p53-PHF20, and reduced p53-MDM2, interaction Pre-vs Post-HVT.

A second important event disrupting the p53-MDM2 interaction and promoting p53 stability is phosphorylation of p53 at serine 15 (68). Pre-HVT, and consistent with this notion, both nuclear and cytosolic p-p53^Ser15^ increased in parallel with the increase in p53 protein content, as previously reported (24), suggesting that phosphorylation of p53 at serine 15 may indeed be involved in the regulation of the p53 protein stability during exercise in human skeletal muscle. In contrast, we report for the first time that there were no exercise-induced changes in p-p53^Ser15^ in either the nuclear or cytosolic fractions after a period of training (i.e., Post-HVT); the increase in resting p-p53^Ser15^ Post-HVT may be a contributing factor for the lack of exercise-induced changes in p-p53^Ser15^ after 40 sessions of HIIT.

This research adds novel information regarding the early molecular events regulating the exercise-induced mitochondrial adaptations in subcellular fractions and how these are altered by an exercise training intervention. We provide evidence that 40 sessions of HIIT blunted the exercise-induced increases recorded pre-training in both nuclear and cytosolic-enriched subcellular fractions, in all of the molecular parameters measured. Although training has previously been shown to blunt some of the exercise-induced adaptations in whole muscle lysates (55), this is the first study to report training-induced blunting of protein changes in the nucleus, where the majority of transcriptional activity takes place, and where an early increase in PGC-1α protein content has been reported to constitute the initial phase of the exercise-induced adaptive response (73). Future studies should investigate if the loss (or reduction) of the exercise-induced increases in markers of mitochondrial adaptations post-training relates solely to the decrease in relative exercise intensity, and/or if this is exacerbated by the continuous repetition of the same exercise stimulus during the training intervention. Well-designed experiments comparing exercise sessions repeated pre- and post-training at the same relative exercise intensity and at different time points during the training intervention (even after only 1 or 2 training sessions to determine the role, if any, of the “first bout effect”) should provide valuable insight into the mechanisms driving this phenomenon.

## Acknowledgements

We thank the participants for their time, effort and commitment to this study. The authors would like to acknowledge Ms. Elise Brentnall and Mr. Maarten Missinne for their valuable help in data collection and biochemical analyses, respectively.

## Author contributions

D. J. Bishop and C. Granata designed the research; C. Granata and R. S. F. Oliveira conducted the research; C. Granata, R. S. F. Oliveira, J. P. Little, and D. J. Bishop analyzed and interpreted the data; C. Granata wrote the manuscript; C. Granata, R. S. F. Oliveira, J. P. Little, and D. J. Bishop critically revised and contributed to the manuscript; C. Granata and D. J. Bishop have primary responsibility for final content. Data collection took place at Victoria University. Muscle analysis took place at Victoria University and the University of British Columbia Okanagan. All persons designated as authors qualify for authorship, and all those qualifying for authorship are listed. All authors have read and approved the final manuscript.

## References

1. Alberts B, Johnson A, Lewis J, Raff M, Roberts K, and Walter P. Molecular Biology of the Cell. 5th edition. New York, NY: Garland Science, 2007.

2. Andersen CL, Jensen JL, and Ørntoft TF. Normalization of Real-Time Quantitative Reverse Transcription-PCR Data: A Model-Based Variance Estimation Approach to Identify Genes Suited for Normalization, Applied to Bladder and Colon Cancer Data Sets. Cancer Res 64: 5245–5250, 2004.

3. Anderson RM, Barger JL, Edwards MG, Braun KH, O’Connor CE, Prolla TA, and Weindruch R. Dynamic regulation of PGC-1α localization and turnover implicates mitochondrial adaptation in calorie restriction and the stress response. Aging Cell 7: 101–111, 2008.

4. Bishop DJ, Botella J, Genders AJ, Lee MJ-C, Saner NJ, Kuang J, Yan X, and Granata C. High-Intensity Exercise and Mitochondrial Biogenesis: Current Controversies and Future Research Directions. Physiology 34: 56–70, 2019.

5. Bishop DJ, Jenkins DG, McEniery M, and Carey MF. Relationship between plasma lactate parameters and muscle characteristics in female cyclists. Med Sci Sports Exerc 32: 1088–1093, 2000.

6. Booth FW, Gordon SE, Carlson CJ, and Hamilton MT. Waging war on modern chronic diseases: primary prevention through exercise biology. J Appl Physiol 88: 774–787, 2000.

7. Canto C, and Auwerx J. PGC-1α, SIRT1 and AMPK, an energy sensing network that controls energy expenditure. Curr Opin Lipidol 20: 98–105, 2009.

8. Cartoni R, Léger B, Hock MB, Praz M, Crettenand A, Pich S, Ziltener JL, Luthi F, Dériaz O, Zorzano A, Gobelet C, Kralli A, and Russell AP. Mitofusins 1/2 and ERRα expression are increased in human skeletal muscle after physical exercise. J Physiol 567: 349–358, 2005.

9. Chen Z-P, McConell GK, Michell BJ, Snow RJ, Canny BJ, and Kemp BE. AMPK signaling in contracting human skeletal muscle: acetyl-CoA carboxylase and NO synthase phosphorylation. American Journal of Physiology-Endocrinology And Metabolism 279: E1202–E1206, 2000.

10. Chen ZP, Stephens TJ, Murthy S, Canny BJ, Hargreaves M, Witters LA, Kemp BE, and McConell GK. Effect of exercise intensity on skeletal muscle AMPK signaling in humans. Diabetes 52: 2205–2212, 2003.

11. Cui G, Park S, Badeaux AI, Kim D, Lee J, Thompson JR, Yan F, Kaneko S, Yuan Z, Botuyan MV, Bedford MT, Cheng JQ, and Mer G. PHF20 is an effector protein of p53 double lysine methylation that stabilizes and activates p53. Nature Structural and Molecular Biology 19: 916–924, 2012.

12. De Filippis E, Alvarez G, Berria R, Cusi K, Everman S, Meyer C, and Mandarino LJ. Insulin-resistant muscle is exercise resistant: Evidence for reduced response of nuclear-encoded mitochondrial genes to exercise. Am J Physiol Endocrinol Metab 294: E607–E614, 2008.

13. Dumke CL, Davis JM, Murphy EA, Nieman DC, Carmichael MD, Quindry JC, Triplett NT, Utter AC, Gross Gowin SJ, Henson DA, McAnulty SR, and McAnulty LS. Successive bouts of cycling stimulates genes associated with mitochondrial biogenesis. Eur J Appl Physiol 107: 419–427, 2009.

14. Eaton M, Granata C, Barry J, Safdar A, Bishop D, and Little JP. Impact of a single bout of high-intensity interval exercise and short-term interval training on interleukin-6, FNDC5, and METRNL mRNA expression in human skeletal muscle. Journal of Sport and Health Science 7: 191–196, 2018.

15. Edgett BA, Foster WS, Hankinson PB, Simpson CA, Little JP, Graham RB, and Gurd BJ. Dissociation of increases in PGC-1α and its regulators from exercise intensity and muscle activation following acute exercise. PLoS One 8: 2013.

16. Egan B, O’Connor PL, Zierath JR, and O’Gorman DJ. Time course analysis reveals gene-specific transcript and protein kinetics of adaptation to short-term aerobic exercise training in human skeletal muscle. PLoS One 8: 2013.

17. Egan B, and Zierath JR. Exercise metabolism and the molecular regulation of skeletal muscle adaptation. Cell Metab 17: 162–184, 2013.

18. Gerhart-Hines Z, Rodgers JT, Bare O, Lerin C, Kim SH, Mostoslavsky R, Alt FW, Wu Z, and Puigserver P. Metabolic control of muscle mitochondrial function and fatty acid oxidation through SIRT1/PGC-1α. EMBO J 26: 1913–1923, 2007.

19. Gilde A, and Van Bilsen M. Peroxisome proliferator-activated receptors (PPARS): regulators of gene expression in heart and skeletal muscle. Acta Physiol Scand 178: 425–434, 2003.

20. Gottifredi V, and Prives C. Getting p53 out of the nucleus. Science 292: 1851–1852, 2001.

21. Granata C, Jamnick NA, and Bishop DJ. Principles of exercise prescription, and how they influence exercise-induced changes of transcription factors and other regulators of mitochondrial biogenesis. Sports Med 48: 1541–1559, 2018.

22. Granata C, Jamnick NA, and Bishop DJ. Training-induced changes in mitochondrial content and respiratory function in human skeletal muscle. Sports Med 48: 1809–1828, 2018.

23. Granata C, Oliveira RSF, Little JP, Renner K, and Bishop DJ. Mitochondrial adaptations to high-volume exercise training are rapidly reversed after a reduction in training volume in human skeletal muscle. FASEB J 30: 3413–3423, 2016.

24. Granata C, Oliveira RSF, Little JP, Renner K, and Bishop DJ. Sprint-interval but not continuous exercise increases PGC-1α protein content and p53 phosphorylation in nuclear fractions of human skeletal muscle. Sci Rep 7: 44227, 2017.

25. Granata C, Oliveira RSF, Little JP, Renner K, and Bishop DJ. Training intensity modulates changes in PGC-1α and p53 protein content and mitochondrial respiration, but not markers of mitochondrial content in human skeletal muscle. FASEB J 30: 959–970, 2016.

26. Groennebaek T, Jespersen NR, Jakobsgaard J, Sieljacks P, Wang J, Rindom E, Musci R, Bøtker HE, Hamilton KL, and Miller BF. Skeletal muscle mitochondrial protein synthesis and respiration increase with low-load blood flow restricted as well as high-load resistance training. Front Physiol 9: 1796, 2018.

27. Gurd BJ, Yoshida Y, McFarlan JT, Holloway GP, Moyes CD, Heigenhauser GJF, Spriet L, and Bonen A. Nuclear SIRT1 activity, but not protein content, regulates mitochondrial biogenesis in rat and human skeletal muscle. Am J Physiol Regul Integr Comp Physiol 301: R67–R75, 2011.

28. Halson SL, Bridge MW, Meeusen R, Busschaert B, Gleeson M, Jones DA, and Jeukendrup AE. Time course of performance changes and fatigue markers during intensified training in trained cyclists. J Appl Physiol 93: 947–956, 2002.

29. Hammond KM, Impey SG, Currell K, Mitchell N, Shepherd SO, Jeromson S, Hawley JA, Close GL, Hamilton LD, Sharples AP, and Morton JP. Postexercise high-fat feeding suppresses p70S6K1 activity in human skeletal muscle. Med Sci Sports Exerc 48: 2108–2117, 2016.

30. Handschin C, Rhee J, Lin J, Tarr PT, and Spiegelman BM. An autoregulatory loop controls peroxisome proliferator-activated receptor γ coactivator 1α expression in muscle. Proc Natl Acad Sci U S A 100: 7111–7116, 2003.

31. Handschin C, and Spiegelman BM. Peroxisome proliferator-activated receptor γ coactivator 1 coactivators, energy homeostasis, and metabolism. Endocr Rev 27: 728–735, 2006.

32. Haupt Y, Maya R, Kazaz A, and Oren M. Mdm2 promotes the rapid degradation of p53. Nature 387: 296–299, 1997.

33. Heesch MW, Shute RJ, Kreiling JL, and Slivka DR. Transcriptional control, but not subcellular location, of PGC-1α is altered following exercise in a hot environment. J Appl Physiol 121: 741–749, 2016.

34. Holloszy JO. Biochemical adaptations in muscle. Effects of exercise on mitochondrial oxygen uptake and respiratory enzyme activity in skeletal muscle. J Biol Chem 242: 2278–2282, 1967.

35. Hood DA. Mechanisms of exercise-induced mitochondrial biogenesis in skeletal muscle. Appl Physiol Nutr Metab 34: 465–472, 2009.

36. Irrcher I, Ljubicic V, Kirwan AF, and Hood DA. AMP-activated protein kinase-regulated activation of the PGC-1α promoter in skeletal muscle cells. PLoS One 3: 2008.

37. Jacobs RA, and Lundby C. Mitochondria express enhanced quality as well as quantity in association with aerobic fitness across recreationally active individuals up to elite athletes. J Appl Physiol 114: 344–350, 2013.

38. Jäger S, Handschin C, St-Pierre J, and Spiegelman BM. AMP-activated protein kinase (AMPK) action in skeletal muscle via direct phosphorylation of PGC-1α. Proc Natl Acad Sci USA 104: 12017–12022, 2007.

39. Jones RG, Plas DR, Kubek S, Buzzai M, Mu J, Xu Y, Birnbaum MJ, and Thompson CB. AMP-activated protein kinase induces a p53-dependent metabolic checkpoint. Mol Cell 18: 283–293, 2005.

40. Kang HM, Ahn SH, Choi P, Ko Y-A, Han SH, Chinga F, Park ASD, Tao J, Sharma K, and Pullman J. Defective fatty acid oxidation in renal tubular epithelial cells has a key role in kidney fibrosis development. Nat Med 21: 37, 2015.

41. Kuang J, Yan X, Genders AJ, Granata C, and Bishop DJ. An overview of technical considerations when using quantitative real-time PCR analysis of gene expression in human exercise research. PLoS One 13: e0196438, 2018.

42. Laursen PB, and Jenkins DG. The scientific basis for high-intensity interval training: Optimising training programmes and maximising performance in highly trained endurance athletes. Sports Med 32: 53–73, 2002.

43. Little JP, Safdar A, Bishop D, Tarnopolsky MA, and Gibala MJ. An acute bout of high-intensity interval training increases the nuclear abundance of PGC-1alpha and activates mitochondrial biogenesis in human skeletal muscle. Am J Physiol Regul Integr Comp Physiol 300: R1303–1310, 2011.

44. Little JP, Safdar A, Cermak N, Tarnopolsky MA, and Gibala MJ. Acute endurance exercise increases the nuclear abundance of PGC-1α in trained human skeletal muscle. Am J Physiol Endocrinol Metab 298: R912–R917, 2010.

45. Londeree BR. Effect of training on lactate/ventilatory thresholds: A meta-analysis. Med Sci Sports Exerc 29: 837–843, 1997.

46. Luft R. The development of mitochondrial medicine. Proc Natl Acad Sci USA 91: 8731–8738, 1994.

47. Marchenko ND, Hanel W, Li D, Becker K, Reich N, and Moll UM. Stress-mediated nuclear stabilization of p53 is regulated by ubiquitination and importin-α3 binding. Cell Death Differ 17: 255–267, 2010.

48. McConell GK, Lee-Young RS, Chen ZP, Stepto NK, Huynh NN, Stephens TJ, Canny BJ, and Kemp BE. Short-term exercise training in humans reduces AM PK signalling during prolonged exercise independent of muscle glycogen. The Journal of physiology 568: 665–676, 2005.

49. Morrison D, Hughes J, Della Gatta PA, Mason S, Lamon S, Russell AP, and Wadley GD. Vitamin C and E supplementation prevents some of the cellular adaptations to endurance-training in humans. Free Radic Biol Med 89: 852–862, 2015.

50. Murton AJ, Billeter R, Stephens FB, Des Etages SG, Graber F, Hill RJ, Marimuthu K, and Greenhaff PL. Transient transcriptional events in human skeletal muscle at the outset of concentric resistance exercise training. J Appl Physiol 116: 113–125, 2013.

51. Nordsborg NB, Lundby C, Leick L, and Pilegaard H. Relative workload determines exercise-induced increases in PGC-1α mRNA. Med Sci Sports Exerc 42: 1477–1484, 2010.

52. Nunnari J, and Suomalainen A. Mitochondria: in sickness and in health. Cell 148: 1145–1159, 2012.

53. Oren M. Regulation of the p53 tumor suppressor protein. J Biol Chem 274: 36031–36034, 1999.

54. Park S, Kim D, Dan HC, Chen H, Testa JR, and Cheng JQ. Identification of Akt interaction protein PHF20/TZP that transcriptionally regulates p53. J Biol Chem 287: 11151–11163, 2012.

55. Perry CGR, Lally J, Holloway GP, Heigenhauser GJF, Bonen A, and Spriet LL. Repeated transient mRNA bursts precede increases in transcriptional and mitochondrial proteins during training in human skeletal muscle. J Physiol 588: 4795–4810, 2010.

56. Pfaffl MW, Tichopad A, Prgomet C, and Neuvians TP. Determination of stable housekeeping genes, differentially regulated target genes and sample integrity: BestKeeper - Excel-based tool using pair-wise correlations. Biotechnol Lett 26: 509–515, 2004.

57. Philp A, Chen A, Lan D, Meyer GA, Murphy AN, Knapp AE, Olfert IM, McCurdy CE, Marcotte GR, Hogan MC, Baar K, and Schenk S. Sirtuin 1 (SIRT1) deacetylase activity is not required for mitochondrial biogenesis or peroxisome proliferator-activated receptor-γ coactivator-1α (PGC-1α) deacetylation following endurance exercise. J Biol Chem 286: 30561–30570, 2011.

58. Pilegaard H, Saltin B, and Neufer DP. Exercise induces transient transcriptional activation of the PGC-1α gene in human skeletal muscle. J Physiol 546: 851–858, 2003.

59. Popov D, Zinovkin R, Karger E, Tarasova O, and Vinogradova O. Effects of continuous and intermittent aerobic exercise upon mRNA expression of metabolic genes in human skeletal muscle. J Sports Med Phys Fitness 54: 362–369, 2014.

60. Popov DV, Zinovkin RA, Karger EM, Tarasova OS, and Vinogradova OL. The effect of aerobic exercise on the expression of genes in skeletal muscles of trained and untrained men. Hum Physiol 39: 190–195, 2013.

61. Puigserver P, Rhee J, Lin J, Wu Z, Yoon JC, Zhang CY, Krauss S, Mootha VK, Lowell BB, and Spiegelman BM. Cytokine Stimulation of Energy Expenditure through p38 MAP Kinase Activation of PPARγ Coactivator-1. Mol Cell 8: 971–982, 2001.

62. Puigserver P, and Spiegelman BM. Peroxisome proliferator-activated receptor-γ coactivator 1α (PGC-1α): Transcriptional coactivator and metabolic regulator. Endocr Rev 24: 78–90, 2003.

63. Rodgers JT, Lerin C, Gerhart-Hines Z, and Puigserver P. Metabolic adaptations through the PGC-1α and SIRT1 pathways. FEBS Lett 582: 46–53, 2008.

64. Saleem A, Carter HN, Iqbal S, and Hood DA. Role of p53 within the regulatory network controlling muscle mitochondrial biogenesis. Exerc Sport Sci Rev 39: 199–205, 2011.

65. Scarpulla RC. Nuclear activators and coactivators in mammalian mitochondrial biogenesis. Biochim Biophys Acta, Gene Struct Expression 1576: 1–14, 2002.

66. Scarpulla RC. Transcriptional paradigms in mammalian mitochondrial biogenesis and function. Physiol Rev 88: 611–638, 2008.

67. She QB, Bode AM, Ma WY, Chen NY, and Dong Z. Resveratrol-induced activation of p53 and apoptosis is mediated by extracellular-signal-regulated protein kinases and p38 kinase. Cancer Res 61: 1604–1610, 2001.

68. Shieh SY, Ikeda M, Taya Y, and Prives C. DNA damage-induced phosphorylation of p53 alleviates inhibition by MDM2. Cell 91: 325–334, 1997.

69. Stepto NK, Benziane B, Wadley GD, Chibalin AV, Canny BJ, Eynon N, and McConell GK. Short-term intensified cycle training alters acute and chronic responses of PGC1α and cytochrome c oxidase IV to exercise in human skeletal muscle. PLoS One 7: 2012.

70. Tachtsis B, Smiles W, Lane S, Hawley J, and Camera DM. Acute endurance exercises induces nuclear p53 abundance in human skeletal muscle. Front Physiol 7: 2016.

71. Welinder C, and Ekblad L. Coomassie staining as loading control in Western blot analysis. J Proteome Res 10: 1416–1419, 2011.

72. Wilkinson DJ, Franchi MV, Brook MS, Narici MV, Williams JP, Mitchell WK, Szewczyk NJ, Greenhaff PL, Atherton PJ, and Smith K. A validation of the application of D2O stable isotope tracer techniques for monitoring day-to-day changes in muscle protein sub-fraction synthesis in humans. American Journal of Physiology-Heart and Circulatory Physiology 2014.

73. Wright DC, Han DH, Garcia-Roves PM, Geiger PC, Jones TE, and Holloszy JO. Exercise-induced mitochondrial biogenesis begins before the increase in muscle PGC-1α expression. J Biol Chem 282: 194–199, 2007.

74. Wu Z, Puigserver P, Andersson U, Zhang C, Adelmant G, Mootha V, Troy A, Cinti S, Lowell B, Scarpulla RC, and Spiegelman BM. Mechanisms controlling mitochondrial biogenesis and respiration through the thermogenic coactivator PGC-1. Cell 98: 115–124, 1999.

